# Track-To-Learn: A general framework for tractography with deep reinforcement learning

**DOI:** 10.1101/2020.11.16.385229

**Authors:** Antoine Théberge, Christian Desrosiers, Maxime Descoteaux, Pierre-Marc Jodoin

## Abstract

Diffusion MRI tractography is currently the only non-invasive tool able to assess the white-matter structural connectivity of a brain. Since its inception, it has been widely documented that tractography is prone to producing erroneous tracks while missing true positive connections. Anatomical priors have been conceived and implemented in classical algorithms to try and tackle these issues, yet problems still remain and the conception and validation of these priors is very challenging. Recently, supervised learning algorithms have been proposed to learn the tracking procedure implicitly from data, without relying on anatomical priors. However, these methods rely on labelled data that is very hard to obtain. To remove the need for such data but still leverage the expressiveness of neural networks, we introduce *Track-To-Learn*: A general framework to pose tractography as a deep reinforcement learning problem. Deep reinforcement learning is a type of machine learning that does not depend on ground-truth data but rather on the concept of “reward”. We implement and train algorithms to maximize returns from a reward function based on the alignment of streamlines with principal directions extracted from diffusion data. We show that competitive results can be obtained on known data and that the algorithms are able to generalize far better to new, unseen data, than prior machine learning-based tractography algorithms. To the best of our knowledge, this is the first successful use of deep reinforcement learning for tractography.

## 1 Introduction

Tractography is the process of inferring white-matter structure from diffusion magnetic resonance imaging (dMRI), by using the diffusion signal to model the underlying fibre populations. From this information and initial seed points, the shape of white-matter tissue is iteratively reconstructed following local orientation information [5]. Directions can be chosen deterministically, by always selecting the principal reconstructed direction, or probabilistically by using the diffusion model as a probability density function for the next direction. Alternatively, global tractography algorithms try to iteratively infer all directions at once while minimizing a loss function [26]. However, the method for choosing the next tracking direction is not the only choice one might face, as for example, the method for modelling the diffusion signal will have a high impact on the produced tractograms (i.e., the product of running a tractography algorithm) [23]

Over the years, a plethora of tractography algorithms have been proposed in the hope of enhancing the coverage of the white-matter volume by streamlines without generating too many false-positives (i.e., streamlines connecting regions that should not be connected). Some methods allow the tractography process to sample several possible directions and select the most promising one [18], others retro-actively modify the fiber orientation diffusion functions (fODF) to redirect the tracking directions [41] and others inject anatomical priors based on atlases on a per-bundle basis[61].

Despite all these efforts, several issues remain as tractography is an *ill-posed* problem that tries to infer global connectivity only from local information [29]. Emerging from this observation is the “bottleneck” problem [40]: in areas where several white-matter bundles have to pass through the same “choke point” and then separate, tractography is typically unable to disentangle where to go after exiting the bottleneck. However, this problem does not come from modelling or directional decision making; it stems from the fact that from a local point of view, every “exit” option makes as much sense as the others, resulting in a several false-positive bundles being generated and true-positives being omitted.

Recently, supervised machine learning methods have been proposed in the hope of learning the tracking procedure without the need of strong, hard-to-implement anatomical priors, as well as removing the need for diffusion modelling [36]. Harnessing the pattern matching abilities of machine learning, these methods propose to learn how to infer the next tracking direction directly from raw diffusion data and labelled data in the form of “ground-truth” tractograms.

Generating these ground-truth tractograms is no easy feat. Even for hardened neuroimagery experts, inferring streamline orientation purely from a visual inspection of diffusion imaging is virtually impossible. One way to proceed is to build “phantoms”, devices made of synthetic materials to mimic physical properties of the human brain, like the FiberCup [14,37,38]. Generating ground-truth bundles from these phantoms is doable as the synthetic bundle configuration is visible and tangible. However, being only ever so similar to real human brains, phantoms cannot fully confirm the capabilities of tractography algorithms. An alternative is to generate the most complete tractogram possible using a family of tractography algorithms and to try to reconstruct all true-positive connections with no regards for false-positives. Then, the overly-full tractogram can be virtually dissected and segmented by experts and/or bundle segmentation algorithms [28]. However, even when done by highly-trained experts following a strict protocol, a high inter-observer variability is observed [39,43]. As such, very few datasets with ground-truth streamlines are publicly available today [42].

Therefore, while supervised machine learning offers an appealing alternative to classical tractography algorithms, the very nature of diffusion MRI and tractography calls for methods that do not rely on ground-truth data.

Tangentially, deep reinforcement learning is a form of machine learning that does not require explicit labelled data, and has recently shown promising results in various fields, from robotic [44] to game playing [30,45,55]. Reinforcement learning formalizes the notion of trial and error by letting a learning agent experiment in its environment, providing feedback through a reward signal.

In this work, we propose to use deep reinforcement learning to leverage the power of deep learning for tractography without the need for biased or hard-to-obtain ground-truth data. Our main contributions are as follow:

1. We propose *Track-to-Learn*, a general framework to pose tractography as a reinforcement learning problem.
2. We analyze several components of the proposed framework to make light on their impact on the reconstructed tractograms.
3. We demonstrate competitive results when compared to supervised machine learning and classical tractography algorithms.
4. We demonstrate far superior generalization capabilities to new, unseen datasets compared to prior machine learning methods.

To our knowledge, we are the first to successfully use deep reinforcement learning for tractography.^2^

### 1.1 Related Work

Over the years, a few machine learning methods have been proposed to predict streamline directions directly from data. We review some of them in this section but for a more complete survey, we recommend the paper by Poulin et al.[36].

The first machine learning method used for tractography is the one by Neher et al.[31,32] which employs a Random Forest classifier to decide on the next tracking step to make. The classifier performs a majority vote on 100 directions sampled from a half-sphere in front of the streamline, with a 101-th class added to model the termination of the streamline. As input, the classifier is given the raw diffusion signal at the head of the streamline, resampled to 100 directions, *N* signal samples in front of the streamline, and the last streamline directions followed by the model. As “ground-truth”, streamlines generated from a constrained spherical deconvolution deterministic tracking algorithm (CSD-DET) [49,50] on the ISMRM2015 Tractography challenge dataset [28] were used, as well as the ground-truth streamlines released as part of the dataset. Results were computed on the same dataset, using the Tractometer [9,10] evaluation tool. By comparison to the original ISMRM2015 submissions, the Random Forest was the only one able to reconstruct all 25 bundles from the ground truth as well as to achieve a higher-than-average overlap.

In order to learn streamline tracking more directly from the data, the *Learn-to-Track* algorithm [35] was proposed. The authors argue that streamlines have directional dependencies which go beyond the last streamline segment. To this extent, they use recurrent neural networks (RNN), more specifically Gated-Recurrent Unit [8] networks^3^. As input to their network, the authors also used the raw diffusion signal resampled to 100 directions. Contrary to Neher et al. [31,32], the RNN outputs the new tracking step directly, without having to perform a classification on fixed directions. The method was also trained using tractograms generated by a CSD-DET algorithm, and managed to outperform most of the original ISMRM2015 tractography challenge submissions.

Similarly to *Learn-to-Track*, Benou et al. introduced the *DeepTract* algorithm [6]. The algorithm also uses RNNs but instead outputs an orientation distribution function (ODF) learned from reference tracks, separates the ODF into classes and then performs classification much like Neher et al. [31,32]. This allows them to perform deterministic as well as probabilistic tractography. They report competitive results when compared to the ISMRM2015 Tractography Challenge submissions and other ML-based tractography methods when trained using either ground-truth data or tracks generated from MITK [60].

Analogously, Wegmayr et al. proposed the *iFOD3* [59] method which uses a feed-forward neural network. As with the other methods, the authors use the raw resampled signal. However, to add information about the neighborhood of the head of the streamline, a bloc of 3 × 3 × 3 voxels is fed to the network instead of the signal coming from a single voxel. Also, because feed-forward networks have no temporal memory, the authors added the last four directions as input to their neural network. While the other methods relied on seeding from the white-matter, Wegmayr et al. chose to seed at the white-matter/grey-matter interface. Furthermore, contrary to other methods, the authors chose to generate streamlines from the iFOD2 algorithm [48]. Results from the Tractometer evaluation tool revealed a high valid connection rate and a low number of invalid bundles, but also a very low overlap.

Subsequently, Wegmayr et al. proposed *Entrack* [58]: a probabilistic machine-learning model for tractography. Building on their previous method, the output of the model is a Fischer-von-Mises (FvM) distribution instead of a deterministic output. The algorithm allows for probabilistic tracking and uncertainty quantification. This results in increased robustness, reduced overfitting and generally better performance compared to their previous work. However, the method still is dependent on labelled-data and does not outperform its reference classical method.

Similar to our approach, an RL method using expanding graphs was first proposed to reduce false positives [56]. The method used ground-truth bundles from the ISMRM2015 Tractography Challenge dataset to infer starting points and end goals for the tracking. It works by generating an expanding graph to discover paths linking the starting and goal regions, and learning a value for each node of the graph using Temporal Difference Learning [46]. Then, the optimal path connecting the starting and ending regions can be inferred from the learned value function. Unfortunately, the reward function remains undisclosed. While this method is interesting, it presumes the existence of ground-truth data, or at least starting and ending regions, which are not available in clinical settings. The method also uses positions as input states, which makes the algorithm non-robust to rotation or translation in the diffusion data. Finally, it is unclear if the algorithm is trained on a per-bundle basis or learns a value function for the whole brain.

## 2 Method

### 2.1 Preliminaries

Reinforcement learning is a framework that allows to solve problems via a sophisticated trial and error process, by placing a learning agent in an environment and providing it with a reward signal based on actions performed. The agent then tries to adapt its strategy to maximize its expected cumulative reward. This trial and error process is formalized as an infinite-horizon Markov Decision Process (MDP) defined by the tuple *(S, A, p, r*), where *S* is the space of all possible states and *A* is the space of all possible actions. *p* indicates the transition probability *p*(*s*_*t*+1_|*s*_*t*_, *a*_*t*_), *s*_*t*_, *s*_*t*+1_ *∈ S, a*_*t*_ *∈ A*, the probability of landing in state *s*_*t*+1_ by executing action *a*_*t*_ in state *s*_*t*_ at time *t. r*(*s*_*t*_, *a*_*t*_), usually shortened to just *r*_*t*_, is the reward obtained from the environment by executing action *a*_*t*_ in state *s*_*t*_ at time *t*. The agent will have a policy *π*(*a*|*s*) which denotes the probability of taking action *a* at state *s*.

RL algorithms typically use the following loop: An initial state *s*_0_ is generated by the environment. An action *a*_0_ is then generated according to the agent’s policy *π*(*a*_0_| *s*_0_) and sent back to the environment so that a new state *s*_1_ is computed together with a reward *r*_0_. Afterwards, the agent uses the next state *s*_1_ to produce action *a*_1_, then a new state *s*_2_ and a new reward *r*_1_ are returned by the environment, and so on. The loop continues for a number *T* of timesteps, or until a stopping criterion is met according to the environment. The series of states and actions that occur is called an *episode*. At the end of an episode, the environment is reset to an initial state and a new episode starts.

The overarching objective of RL is to find the optimal policy *π*(*a*|*s*) that maximizes the expected sum of future rewards. In that perspective, it is common to define the *value function, i*.*e*. the expected reward in state *s*_*t*_ and time *t*:

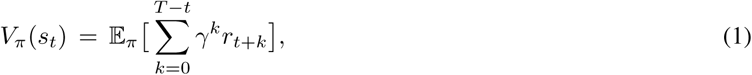

where 0 ≪ *γ* < 1 is a discount factor preventing the cumulative reward to move towards infinity in the case of episodes with infinite length. The discount factor has the property of promoting short-term reward when adjusted towards lower values and long-term reward when adjusted towards higher values. Another commonly-used function is the *state-action value function*, or simply *Q*-function

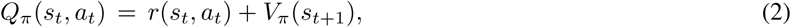

which is the expected upcoming reward that one would get by selecting action *a*_*t*_ at state *s*_*t*_ and then following the policy *π* in subsequent states. To determine which action is best in which state, RL algorithms tend to try to maximize the Bellman Equation defined as

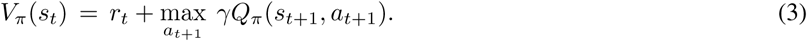

Assuming we have the optimal *Q*-function, that is the state-action value function that produces the true expected return for the optimal policy, then the optimal policy only consists of iteratively selecting the action that maximizes the *Q*-function. Finding this optimal pair of evaluation function and policy is the goal of reinforcement learning.

### 2.2 Proposed framework

We formulate the tractography process as a reinforcement learning problem. The diffusion volume acts as the environment, where a state *s* is the diffusion signal at the head of the streamline. At first, a tracking seed is generated. The diffusion signal *s*_0_ at the seed position is sent to the agent. The agent predicts the tracking direction *a*_0_, which is sent back to the environment to be normalized to a given tracking step size and used to calculate the new position of the streamline’s head. The tracking direction is compared against the local fODF peaks and a reward *r*_0_ is sent back to the agent, along with state *s*_1_. The loop continues until the tracking goes out of a precomputed tracking mask, or the angle between two consecutive segment is too high. The framework is very general and allows for the user to implement each component as they wish. Figure 1 illustrates the proposed framework.

**Figure 1:**
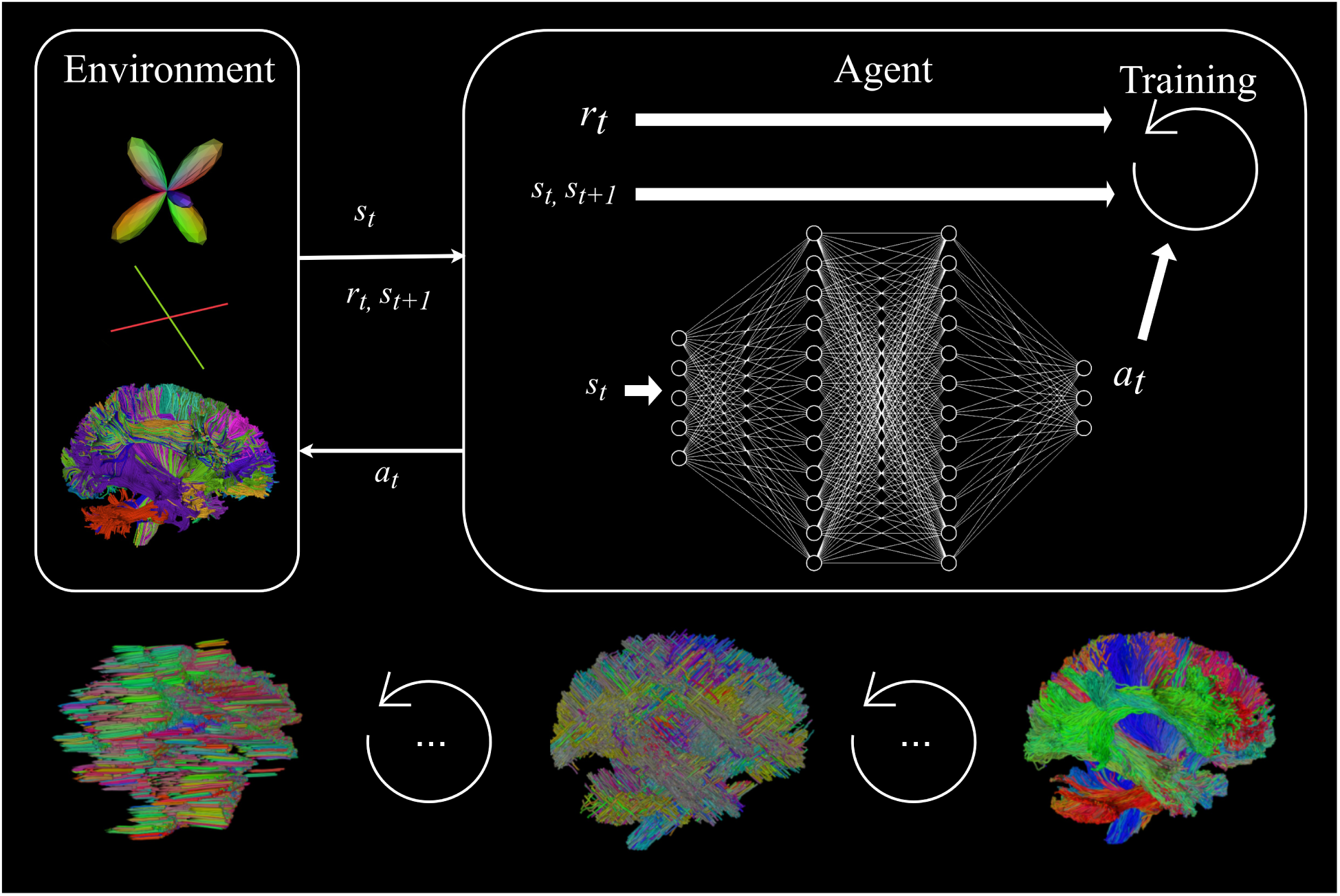
The reinforcement-learning-for-tractography loop as well as the evolution of produced tractograms. Top part: To the left, the environment produces a state *s*_*t*_ from the diffusion signal (see section 2.2.2 for a description) as well as a reward *r*_*t*_, which is computed from the last tracking step and the peaks extracted from the fODFs at the streamline’s head. Both are given to the agent: the state is used as input to the policy (a neural network) to produce an action *a*_*t*_, in this case a new tracking step, and the reward is used to train the agent to produce better actions. The environment receives the action *a*_*t*_, normalizes it to the chosen step size and computes the new position of the streamline’s head. A new state *s*_*t*+1_ and new reward *r*_*t*+1_ is returned and the cycle continues. Bottom part: Tractograms produced by agents before training begins, during training and at the end of training.

#### 2.2.1 Environment

In RL, the role of the environment is to produce initial states, compute new states from actions, handle episode termination, and produce rewards. As such, in our context, the environment produces initial states from seeds generated from a seeding mask. The states correspond to the signal (described in section 2.2.2) at the current streamline’s *head*, or last position. The state is sent to the agent, which produces an action, *i*.*e*. a tridimensional vector which is the next tracking direction. The environment then re-scales the action to a pre-determined tracking step and computes the next streamline’s position, which leads to computing the next state. The reward of the current timestep is also computed, as well as a marker *d*_*t*_ indicating whether or not the tracking for the current streamline is over (cf. section 2.2.3). Streamline termination depends on the following conditions:

1. The streamline exits the white-matter mask;
2. The streamline exceeds a predetermined maximum length;
3. The angle between two consecutive segments in the streamline exceed a maximum angle.
4. The streamline exceeds a maximum cumulative angle between segments.

The angle between streamline segments is defined as

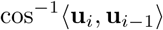

and **u**_*i*_ are normalized streamline segments. If none of these conditions is satisfied, tracking continues and the tuple (*s*_*t*+1_, *r*_*t*_, *d*_*t*_) is returned to the agent.

In itself, tractography is a two-step process. Because diffusion is by nature symmetric, tracking has to first follow one “direction” of the diffusion signal, and then go back to its original position to follow the “other direction”. As such, and distancing ourselves from the standard RL setting, two environments are needed. The first one performs tractography from initial seeds and produces “half-streamlines”. The second (or *reverse*) environment takes in half-streamlines and reverses them so that the initial seed point becomes the head of the streamline. To allow a form of self-supervised learning, tracking is still performed on the reversed streamlines starting from the new initial points, states are still sent to the agent, and rewards are calculated according to the new actions. However, to preserve the integrity of the flipped half-streamlines and to guide the tracking process, actions that are computed by the agent while the agent is retracing its steps are discarded and replaced with the pre-exisiting half-streamlines segments. When the reversed half-streamline has been reconstructed, tracking continues as normal. The process is illustrated in figure 2.

**Figure 2:**
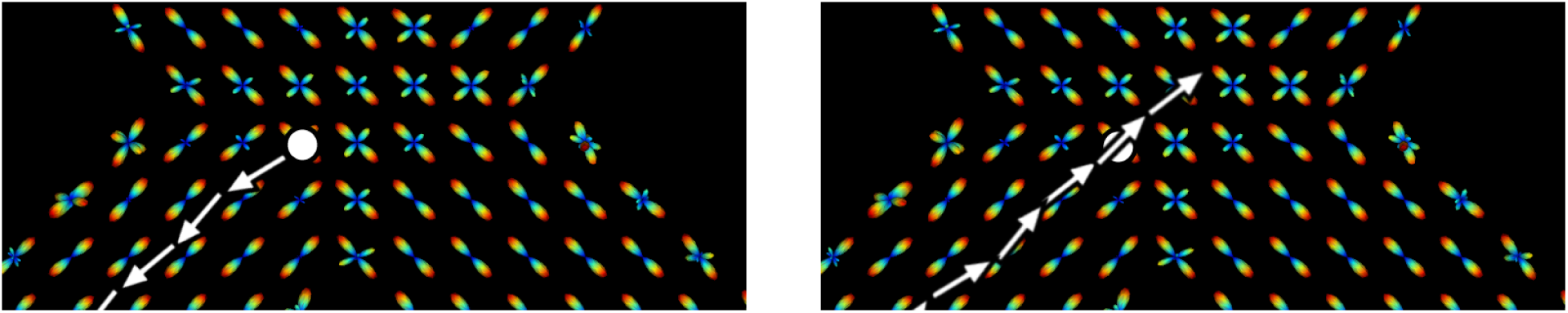
“Forward” and “backward” tracking processes. *Left*: the “forward” environment initiates a seed (white dot) and the tracking starts in one direction, indicated by arrows. Once the tracking is over, the half-streamline is flipped and sent to the backwards environment. *Right*: the half-streamline is flipped, the initial seed point (white dot) becomes the head of the streamline, and the tracking continues.

To take advantage of the batching abilities of neural networks as well as to simulate multiple agents learning at the same time, multiple streamlines are initialized by the “forward” environment and handled at the same time. As such, a list of initial states 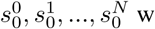 where *N* corresponds to the batch size, is sent to the agent. The agent then answers with a batch of 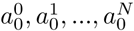 actions which results into a series of *N* new states and *N* rewards and the cycle goes on. As individual streamlines terminate, their corresponding states are removed from the batch and are appended to a tractogram. When all the forward half-streamlines are terminated, the streamlines in the tractogram are flipped and fed to the backward environment.

To align ourselves with most of the recent work done in reinforcement learning, the environment is implemented following the OpenAI Gym[7] specification. The forward environment exposes the *reset* method, which takes in no parameter and returns the initial state *s*_0_, as well as the *step* method, which takes in the action *a*_*t*_ and returns the next state *s*_*t*+1_, reward *r*_*t*_ and completion signal *d*_*t*_. The backward environment follows the same specification, but its *reset* method takes in half-streamlines that have been flipped.

#### 2.2.2 Input signal

The strict Markovian requirements of off-the-shelf RL models does not apply to brain tractography. First, RL algorithms typically assume a Markovian process of order 1, where any state *s*_*t*_ depends only on the state and action *s*_*t*−1_, *a*_*t*−1_ before it. This principle is named the *Markov Property*. This property is not respected when the transition probabilities correspond to *p*(*s*_*t*_ |*s*_*t*−1_, *a*_*t*− 1_, …, *s*_*k*_, *a*_*k*_) with *k* < *t* −1. As illustrated by figure 2 and mentioned by Poulin et al.[35], streamlines have directional priors that go much beyond just local information. As such, the action *a*_*t*_ for propagating the streamline in a certain direction cannot be only driven by the local information contained in *s*_*t*_, but by the last couple of states/actions the system has visited/made before.

Second, RL assumes that state *s*_*t*_ contains all of the information needed to perform the right action *a*_*t*_. Unfortunately, the true state of a brain structure amounts to the sub-pixel cellular structure of the white-matter which is unavailable non-invasively. *In-vivo* tractography relies on diffusion MRI data which amounts to an *observation o*_*t*_ of the *true* state *s*_*t*_. This is much like playing a video game by looking at the screen: the *observations* fed to the agents are the pixels displayed while the *true* state lies in the console registers.

All in all, our proposed model is a Partially-Observable Markov Decision Processes (POMDP) of order greater than 1. POMDPs are defined by the tuple (*S, A, p, r, O*, Ω), where *S* is the set of all true states the environment can have, *A* the set of all actions possible, *p* the transition probabilities as described before, *O* the set of all possible observations and Ω the observation probabilities Ω(*o*_*t*_|*s*_*t*_, *a*_*t*_).

While the observations fed to the agent can be the raw input diffusion signal as in Wegmayr et al.[59] and Poulin et al.[35], we adopted a different approach. As illustrated in figure 3, the input signal fed to our agent are spherical harmonic (SH) coefficients representing fiber ODFs [11,49] obtained by a constrained spherical deconvolution [49]. To add spatial information, the signal from the six immediate neighbours (up, down, left, right, front, back) at the streamline’s end is appended to the input. To add further contextual information, the tracking mask values at these neighbouring points are also appended to the signal. Finally, to alleviate the effects of the directional dependencies on streamlines and inject additional directional information, the last four streamline segments are appended to the input signal, much like Wegmayr et al.[59]. All things considered, we end up with a 215-dimensional input: 28 × 7 SH coefficients + 7 mask values + 4 × 3 vector components for the previous directions.

**Figure 3:**
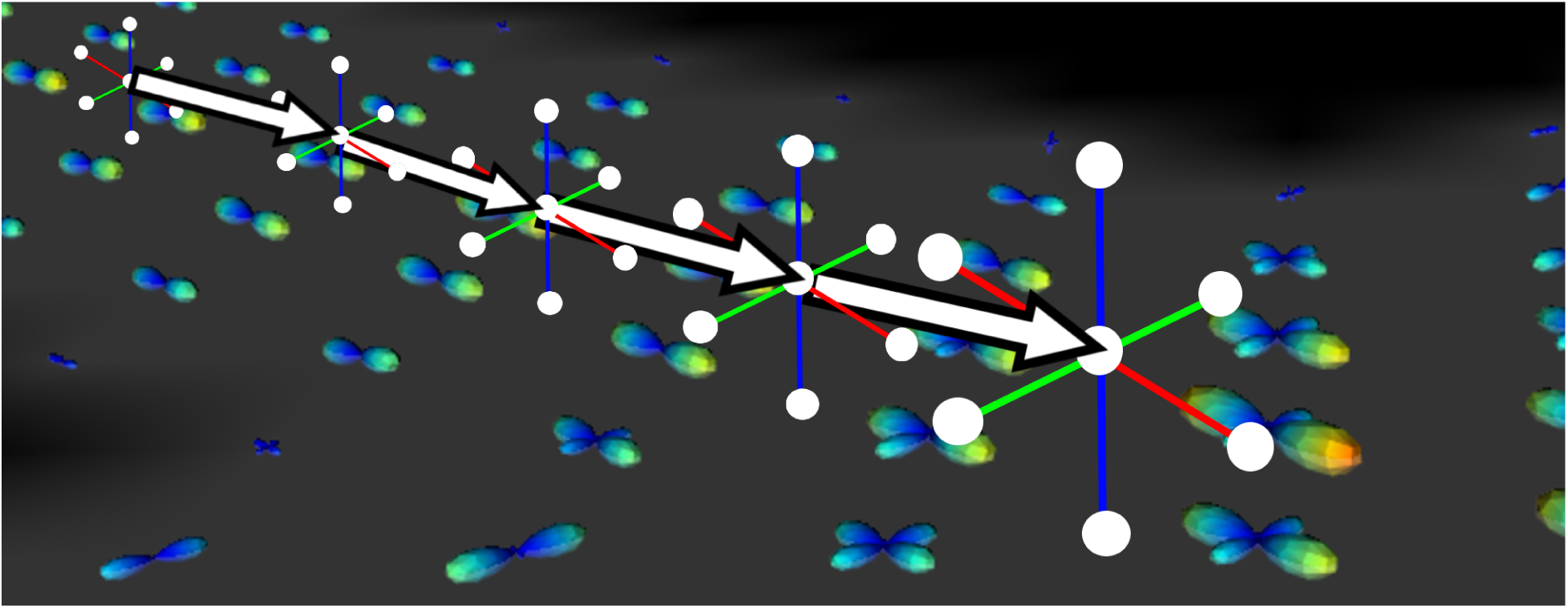
Illustration of the input signal as well as the neighbouring, directional and mask information. SH coefficients are represented by fODF glyphs, the tracking mask is grey where tracking is allowed and black where it is not. Streamline segments are represented by arrows. White circles represent where the input signal is computed.

#### 2.2.3 Reward function

To guide the learning process, a suitable reward function must be provided to the agent. Because the agent is not given any explicit goal, the reward function (and its maximization by the agent) must encompass what the true underlying goal is, or what the desired behaviour of the agent should be. Defining an explicit goal to tractography is unclear. As mentioned before, the exact reconstruction of the ground-truth data should not be an end by itself, as we do not have access to it.

For our implementation of the proposed Track-To-Learn framework, we take a look at the behaviour of classical deterministic tractography algorithms for inspiration: how do they perform tractography? From their local model, they extract peaks and choose the one most aligned with their previous direction, propagate the streamline forward in the peak’s direction and do so until the angle with their previous direction is too high or tracking goes out of the WM mask. Therefore, if we want to learn a tractography algorithm, we should promote behaviours that are executed by one.

We define the reward function for our environment as follows: the reward function is the absolute^4^ cosine distance between the action performed by the agent and the peak most aligned with it, multiplied by the cosine distance between the last streamline segment and the action performed:

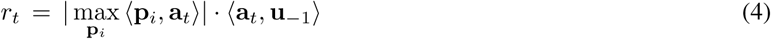

where **p**_*i*_ are the peaks at the head of the streamlines, **a**_*t*_ is the action produced by the agent at time *t*, **u** _−1_ is the last streamline segment before the action is appended to it and **p**_*i*_, **a**_*t*_, **u** _−1_ are all normalized. This allows the agent to produce streamlines that are aligned with the underlying signal while enforcing streamline smoothness. An illustration of the relation between the reward function and streamlines is available in figure 4.

**Figure 4:**
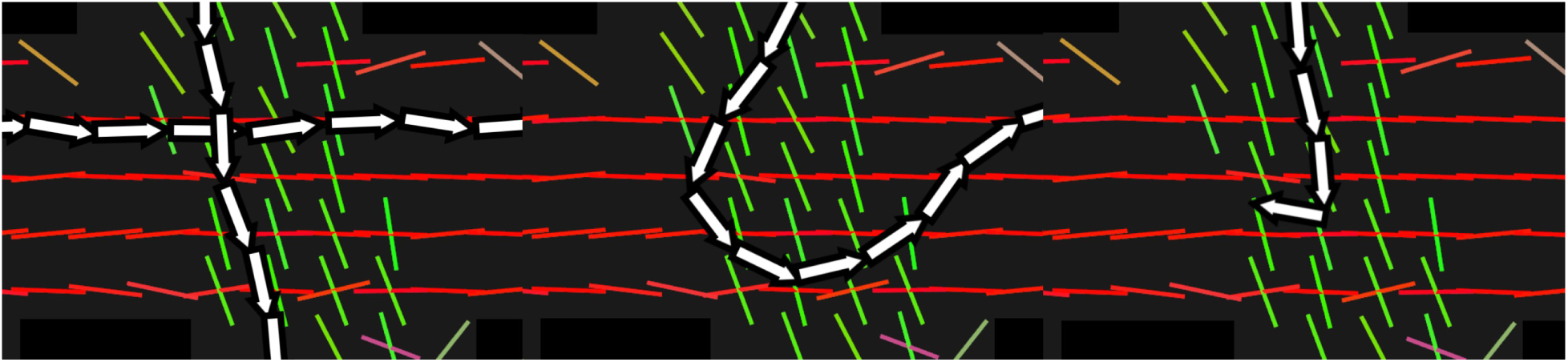
Illustration of the reward signal. The green and red sticks are ODF peaks and the white arrows are streamline segments. *a*) Multiple streamline segments are well aligned with peaks extracted from the fODFs and therefore receive high reward (0 ≪ *r*_*t*_ < 1); *b*) Multiple streamlines segments are badly aligned with the peaks and do not receive a high reward (0 < *r*_*t*_ ≪ 1); *c*) Streamline segments have good alignment with the peaks, however, the last two segments have a high angle between them, bringing the reward at that point closer to −1 and terminating the tracking process.

#### 2.2.4 Agents

To train agents to reconstruct streamlines, we chose two algorithms: Twin-Delayed Deep-Deterministic Policy Gradient (TD3) [15] and and Soft Actor-Critic (SAC) [21]. We selected TD3 for its simplicity and high performance on robotic control tasks. The TD3 algorithm uses the Actor-Critic framework, where a policy network (the actor) performs actions and a critic network infers what the expected value of the actions performed by the actor will be. The “Twin” part in TD3 comes from the fact that it uses *two* critics instead of one to fight overestimation of the state-action values due to noise in the data [22]. All three models are three-layer fully-connected neural networks with a width of 1024 or 2048, depending on the experiment. The actor predicts deterministic actions, a 3D vector indicating the next unnormalized tracking step.

Analogously, we chose the SAC algorithm for its similarity to TD3 with the added benefit of it having a stochastic policy, meaning that the actor predicts the mean and variance of a distribution instead of actions directly. This allows us to remove the need to explicitly control the exploration done by the agent during training (more on this in section 5.5). SAC also employs the Actor-Critic framework and also uses two critics instead of one. Besides the output of the actor, the architecture employed by SAC is the same as TD3. Both algorithms were implemented using the PyTorch [34] framework.

Another reason for these two algorithms are their deterministic (TD3) and stochastic (SAC) nature which will allow us to compare our approach to conventional deterministic and probabilistic tracking methods.

At test time, to promote a better coverage of the white-matter at test time, noise proportional to the FA is added to the actions such that

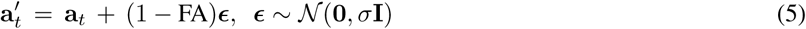

where **a**_*t*_ is the action at timestep *t*, FA is the fractional anisotropy value at the head of the streamline and *σ* is the tracking noise. We chose this strategy instead of applying the same amount of noise everywhere because we presume the algorithm to benefit more from exploratory noise in the more ambiguous, lower FA regions (such as crossings) while lowering the risk of ending the tracking prematurely in simpler, high FA regions because of noise. Even though SAC uses a stochastic policy and therefore could do provide its own noise, we decided to use its policy deterministically at test time and explicitly add noise to better control the noise output by the agent.

Hyperparameters were chosen on a per-experiment basis through a Bayesian hyperparameter search using comet.ml^5^, by selecting the agents reporting the best performance according to the evaluation metrics defined in section 3.2.

## 3 Experimental protocol

### 3.1 Datasets

To train and validate the proposed method, three datasets, both *in silico* and *in vivo*, were used.

#### FiberCup

The FiberCup [14,37,38] is a synthetic dataset that replicates a coronal slice of the brain. The dataset contains 3 axial slices acquired on 30 directions with a b-value of 1000 and an isometric voxel size 3mm. The 7 groundtruth bundles present challenging configurations such as fanning, crossing and kissing fibers. No further preprocessing of the data was necessary as the simulated version of the FiberCup is already free of artifacts. From the data, a volume of SH coefficients, a fractional anisotropy (FA) map and peaks from the fODFs were produced using dipy [17] and scilpy^6^, and a tracking and seeding mask was extracted by thresholding the FA map. Training was performed at 2 seeds per voxel for 100 episodes and testing was performed at 33 seeds per voxel.

#### ISMRM2015

The ISMRM2015 Challenge dataset [28] is a more realistic synthetic dataset. It was generated by tracking on multiple subjects from the Human-Connectome Project dataset [19], and the set of tractograms was manually cleaned and segmented into 25 bundles representing both the major pathways of the white-matter as well as smaller, harder to track bundles. The Fiberfox [33] tool was then used to generate a diffusion-weighted volume as well as a T1 image from the 25 bundles. As such, the dataset does have a proper ground-truth but is still considered synthetic. We chose to preprocess the dataset using Tractoflow [2,12,17,24,27,47,51], which lead to an SH volume of 1× 1× 1mm resolution. The FA and peaks were also computed by Tractoflow, and the tracking and seeding mask was also obtained by thresholding the FA map and then manually cleaning the mask. Training was performed at 1 seeds per voxel for 100 episodes and testing was done at 7 seeds per voxel.

#### Human Connectome Projet

The HCP 1200 dataset [52–54] is a staggering collection of 1200 healthy subjects acquired on a 3T MRI scanner. The diffusion data was acquired at a 1.25mm isometric voxel size for 270 directions distributed over 3 shells. For our project, we used a single subject (ID: 100206) from the HCP 1200 test-retest subset, which was processed with Tractoflow to obtain similar inputs as with the ISMRM2015 dataset. As opposed to the other datasets, the HCP dataset does not have a publicly available set of “ground-truth” streamlines. However, because our method does not rely on labelled data, we could still train on it. Like the ISMRM2015 dataset, training was performed at 1 seeds per voxel for 100 episodes and testing was done at 7 seeds per voxel.

### 3.2 Evaluation metrics

To gauge performance against prior methods, the Tractometer [9,10] tool was used. The Tractometer segments a tractogram into “valid” and “invalid” streamlines according to ground-truth bundles and then extracts performance measures. While it was first used on the FiberCup, it was later on adapted to the ISMRM2015 dataset. The Tractometer reports the true-positive rate (streamlines connecting regions that should be connected) as *valid connections* (VC), the false-positive rate (streamlines connecting two regions that should not be connected) as *invalid connection* (IC) and the true-negative rate (the ratio of streamlines ending prematurely without connecting two regions) as no-connections (NC). The tractometer also reports the number of valid bundles (VB), the number of ground-truth bundles with at least one streamline reconstructed, as well as invalid bundles (IB), the number of regions that are connected by at least one streamline but should not be. To assess the reconstruction of the white-matter volume, the Tractometer also reports mean overlap (OL) and mean overreach (OR), a voxel-wise measure of the overlap (having the same voxels containing streamlines) and overreach (extraneous voxels containing streamlines) between reconstructed and ground-truth bundles.

All tracking was performed using a 0.75 voxel step size and a maximum length of (200 steps*/*step size). We restrict the maximum angle between streamline segments at 60 degrees for the tracking done with TD3 and 30 degrees for the tracking done with SAC. At test time, streamlines under 20mm, over 200mm or with a projected angle higher than 330 degrees are discarded.

### 3.3 Benchmarking & Experiments

Here we present experiments designed to assess the reconstruction and generalization capabilities of our method.

We train our two agents on the FiberCup dataset, and test them on a version of the FiberCup that has been flipped horizontally, aptly dubbed “Flipped”. This allows to measure to which extent our agents can generalize on an environment where the signal distribution is locally different. No ground-truth bundles or masks were used during training. To provide a baseline, we also trained the Learn-to-Track [35] algorithm on the FiberCup dataset with ground truth bundles, both with raw diffusion signal as input, like in the original paper, as well as the fODF SH coefficients and WM mask as input, like for our method. To assess the loss of performance between the training and test datasets, we also report Tractometer metrics for the un-flipped FiberCup. Finally, to provide a more complete comparison, we report metrics from classical deterministic and probabilistic algorithms^7^ which use fODF SH coefficients as inputs (CSD-DET, CSD-PROB). Classical algorithms were executed at 33 seeds per voxels, a step size of 0.75mm, a minimum streamline length of 20mm and a maximum streamline length of 200mm to mirror our experiment protocol.

#### Experiment 2: Performance on the ISMRM2015 dataset

This experiment was designed to mirror state-of-the-art methods by Neher et al.[31,32], Poulin et al.[35] and Benou et al.[6]. We train and test Track-To-Learn agents on the ISMRM2015 dataset and report metrics using the Tractometer tool. However, contrary to prior methods, no ground-truth streamlines or masks were used in the process of training. Comparison is provided against prior supervised learning methods, classical algorithms as well as the mean results of the original ISMRM2015 White-Matter Tractography challenge [28] submissions. Classical methods were executed at 7 seeds per voxels, a step size of 0.75mm, a minimum streamline length of 20mm and a maximum streamline length of 200mm to mirror our experiment protocol.

#### Experiment 3: Generalization through HCP and ISMRM2015 datasets

We again explore the generalization capabilities of the proposed method, however this time on a more complex dataset. As with prior work, we train our agents on the HCP dataset and test it on the ISMRM2015 dataset. To obtain a more complete comparison, we follow the same protocol as experiment 1 with Learn-to-Track [35], but this time train it on the HCP Young Adult dataset [20] using reference tracks from TractSeg [57] and test it on the ISMRM2015 dataset. Subjects 749361, 816653, 814649, 871762, 987983 were used as training set for Learn-to-Track and subjects 704238, 896879 for validation. In further experiments, we provide analysis for some of the proposed method’s hyperparameters.

#### Experiment 4: Impact of the discount parameter

We explore the impact of the discount parameter (*γ*) has on the Tractometer metrics. As mentioned in section 2.1, *γ* prevents the sum of rewards to go towards infinity. From a tractographic point of view, it allows to control how “greedy” the agent can be, or how much importance it gives to rewards that might be produced only far in the future. To analyze the impact of this parameter, we picked our TD3 agent and performed multiple training runs on the FiberCup dataset while having all hyperparameters fixed except *γ*.

#### Experiment 5: Impact of the exploration noise

We analyze the impact of the exploration noise on the Tractometer metrics. Exploration is an important concept in RL: through the process of trial and error, a learning agent has to explore its environment by sometimes trying actions that may not be better than what the agent currently estimates as optimal. However, if an agent explores too much, it might not receive as much reward as it should and the learning could stall. This is aptly named the *Exploration vs. Exploitation* problem in RL.

While SAC handles its exploration on its own through its stochastic policy, TD3 requires explicit noise ϵ ∼ 𝒩 (0, *σ*) to be added to actions performed by the agent during training. As such, the *σ* parameter must be carefully chosen as to allow the agent to discover which actions should be performed at different places in the white-matter, without adding too much noise so that the agent cannot perform optimal actions and reinforce them. We performed multiple training runs on the FiberCup dataset with the TD3 algorithm and fixed all hyperparameters except *σ* to analyze its impact.

#### Experiment 6: Impact of noise at test time

As mentioned in section 2.2.4, noise proportional to the FA is added at test time to promote bundle coverage. As such, we can control how the tracking behaves without having to retrain an agent. We use the Track-To-Learn SAC agent trained for Experiment 2 and perform tracking, without re-training, on the ISMRM2015 dataset with various levels of noise. Because SAC employs a stochastic policy, we can also track using its own probabilistic output instead of artificially adding noise.

## 4 Results

### 4.1 Experiment 1: Generalization through the FiberCup dataset

Table 1 shows competitive performances from our agents when training and testing on the same dataset. Compared to the non-machine learning methods CSD-DET and CSD-PROB, our method achieves better VC and NC rates on the original and flipped dataset at the expense of a slightly worse OL and IC. Note that our reported results outperform all of the submissions from the original Tractometer challenge (cf. [10] for more details).

**Table 1:**
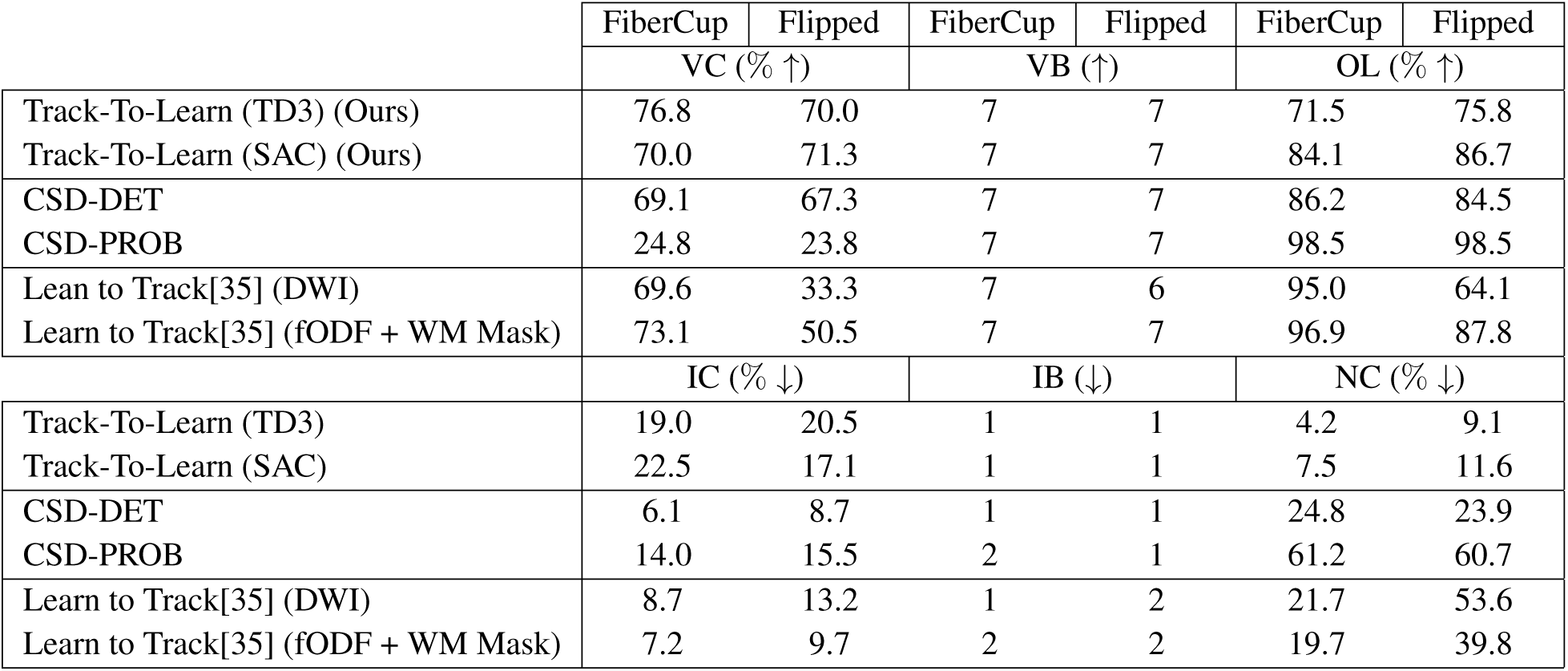
Results obtained by training/testing on the FiberCup. Testing was also done on a flipped version of it.

|  | FiberCup | Flipped | FiberCup | Flipped | FiberCup | Flipped |
| --- | --- | --- | --- | --- | --- | --- |
| | VC (% ↑) | | VB (↑) | | OL (% ↑) | |
| Track-To-Learn (TD3) (Ours) | 76.8 | 70.0 | 7 | 7 | 71.5 | 75.8 |
| Track-To-Learn (SAC) (Ours) | 70.0 | 71.3 | 7 | 7 | 84.1 | 86.7 |
| CSD-DET | 69.1 | 67.3 | 7 | 7 | 86.2 | 84.5 |
| CSD-PROB | 24.8 | 23.8 | 7 | 7 | 98.5 | 98.5 |
| Learn to Track[35] (DWI) | 69.6 | 33.3 | 7 | 6 | 95.0 | 64.1 |
| Learn to Track[35] (fODF + WM Mask) | 73.1 | 50.5 | 7 | 7 | 96.9 | 87.8 |
| | IC (% ↓) | | IB (↓) | | NC (% ↓) | |
| Track-To-Learn (TD3) | 19.0 | 20.5 | 1 | 1 | 4.2 | 9.1 |
| Track-To-Learn (SAC) | 22.5 | 17.1 | 1 | 1 | 7.5 | 11.6 |
| CSD-DET | 6.1 | 8.7 | 1 | 1 | 24.8 | 23.9 |
| CSD-PROB | 14.0 | 15.5 | 2 | 1 | 61.2 | 60.7 |
| Learn to Track[35] (DWI) | 8.7 | 13.2 | 1 | 2 | 21.7 | 53.6 |
| Learn to Track[35] (fODF + WM Mask) | 7.2 | 9.7 | 2 | 2 | 19.7 | 39.8 |

Compared to Learn to track [35], our method reports similar VC rates on the original dataset, but significantly better results on the “Flipped” dataset. While the reported OL is lower than Learn-to-Track on the training dataset, our method has a much better OL on the Flipped dataset. This shows that our method is less prone to overfitting and has better generalization capabilites. Also, although the IC rate is higher for our method, our NC rate is much lower in both settings compared to supervised and classical methods.

Figure 5 provides as visual comparison of the reconstructed valid tracks by Learn-to-Track (DWI) [35] and our SAC agent. We can observe that on the flipped dataset (2nd row) that the reconstruction by Learn-to-Track is missing bundle 1 and that it seems to try to “go back” in bundle 7, possibly indicating some over-fitting. While the reconstruction by the proposed method is a bit “noisier”, we can appreciate that tractograms from the FiberCup and “Flipped” datasets are very similar.

**Figure 5:**
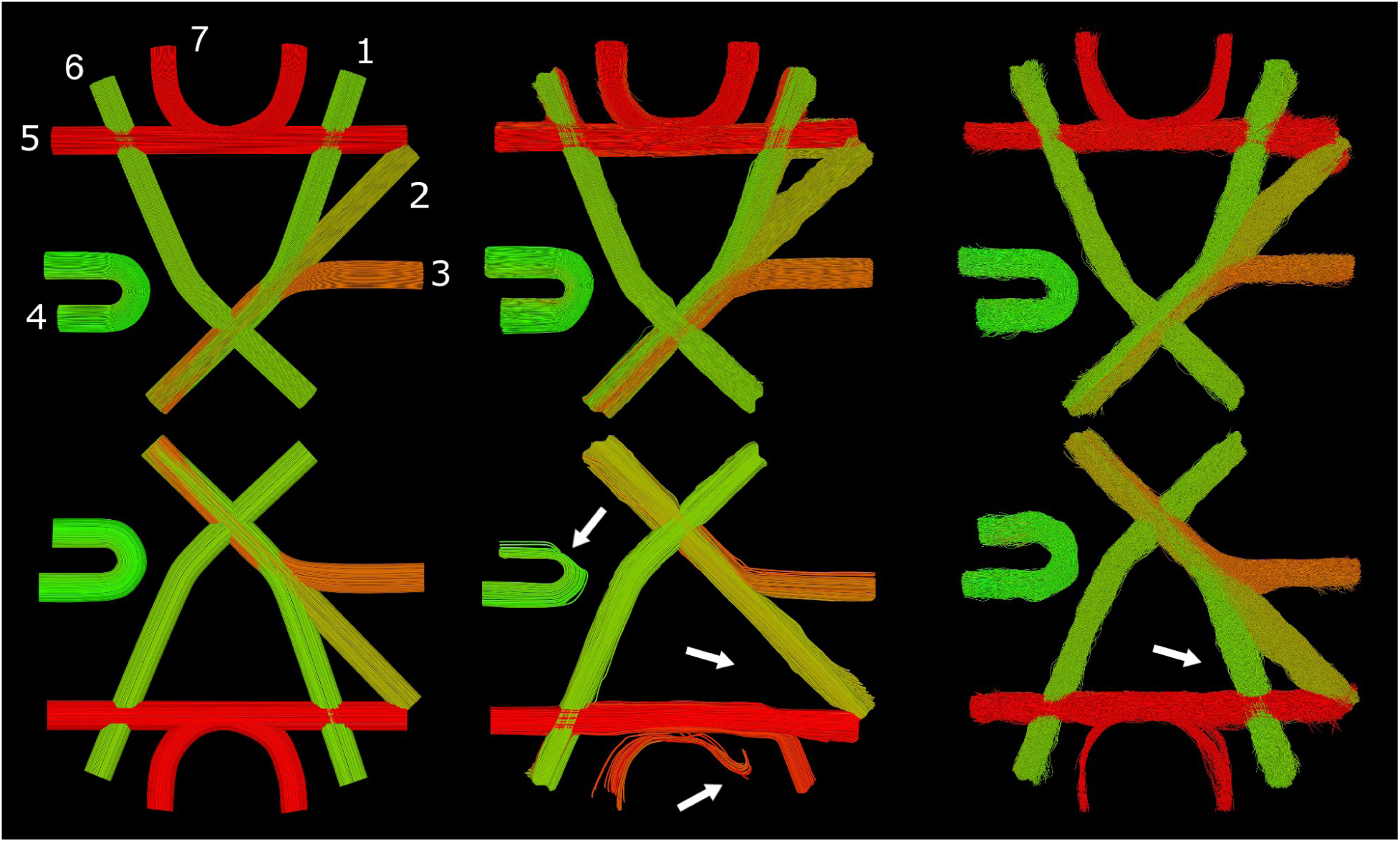
Visualization of the flipped FiberCup dataset and valid streamlines reconstructed by Learn-to-Track [35] and our method. *Left*: Ground-truth fibers with bundle labels. Center: Reconstruction by the Learn-to Track (DWI) algorithm. *Right*: Reconstruction by the SAC agent. Reconstruction of the original (*top*) and flipped (*bottom*) FiberCup dataset. Reconstruction done by the Track-To-Learn agent is quite similar if compared between datasets, while the quality of the reconstruction by the supervised method degrades quite significantly.

### 4.2 Experiment 2: Performance on the ISMRM2015 dataset

Figure 6 provides a visual comparison of the ground-truth bundles against bundles produced by the Track-to-Learn SAC agent. Table 2 shows results of our method trained and tested on the ISMRM2015 dataset. We compare our results against the original ISMRM2015 White-Matter Tractography Challenge submissions, as well as state-of-the-art method by Neher et al. [31,32], Poulin et al.[35] and Benou et al.[6]. Mean results from the ISMRM2015 challenge were extracted from Poulin et al.[35]. Because they only report improvement over mean ISM2015 results and not direct metrics, performances measures for Neher et al. were re-calculated by the authors based on ISMRM2015 mean results.

**Table 2:** Metrics for our method against the mean results for original ISMRM2015 White Matter Challenge sub-missions [28] and prior machine learning methods [6,31,35]. (GT) indicates methods that were trained using the ground-truth streamlines. ‘X’ indicates metrics that were not reported by prior methods.

| | VC (% ↑) | VB (↑) | OL (% ↑) | IC (% ↓) | IB (↓) | NC (% ↓) |
| --- | --- | --- | --- | --- | --- | --- |
| Track-To-Learn (TD3) (Ours) | 55.1 | 23 | 55.8 | 44.2 | 163 | 0.7 |
| Track-To-Learn (SAC) (Ours) | 68.4 | 23 | 62.1 | 30.8 | 161 | 0.7 |
| ISMRM2015 Mean results [28] | 53.6 | 21.4 | 31.0 | 19.7 | 281 | 25.2 |
| CSD-DET | 60.6 | 23 | 51.9 | 33.6 | 134 | 5.8 |
| CSD-PROB | 33.1 | 23 | 79.3 | 53.6 | 178 | 13.3 |
| Neher et al.[31, 32] | 61.1 | 24.3 | 55.1 | X | 86.9 | X |
| DeepTract [6] | 40.5 | 23 | 34.4 | 32.6 | 51 | 22.9 |
| Entrack [58] | 65 | 24 | 60 | X | 117 | X |
| Neher et al.[31, 32] (GT) | 75.6 | 24.3 | 70.6 | X | 86.9 | X |
| DeepTract [6] (GT) | 70.6 | 25.0 | 69.3 | 19.5 | 56 | 9.9 |
| Learn-to-Track [35] (GT) | 41.6 | 23 | 64.4 | 45.6 | 130 | 12.8 |

**Figure 6:**
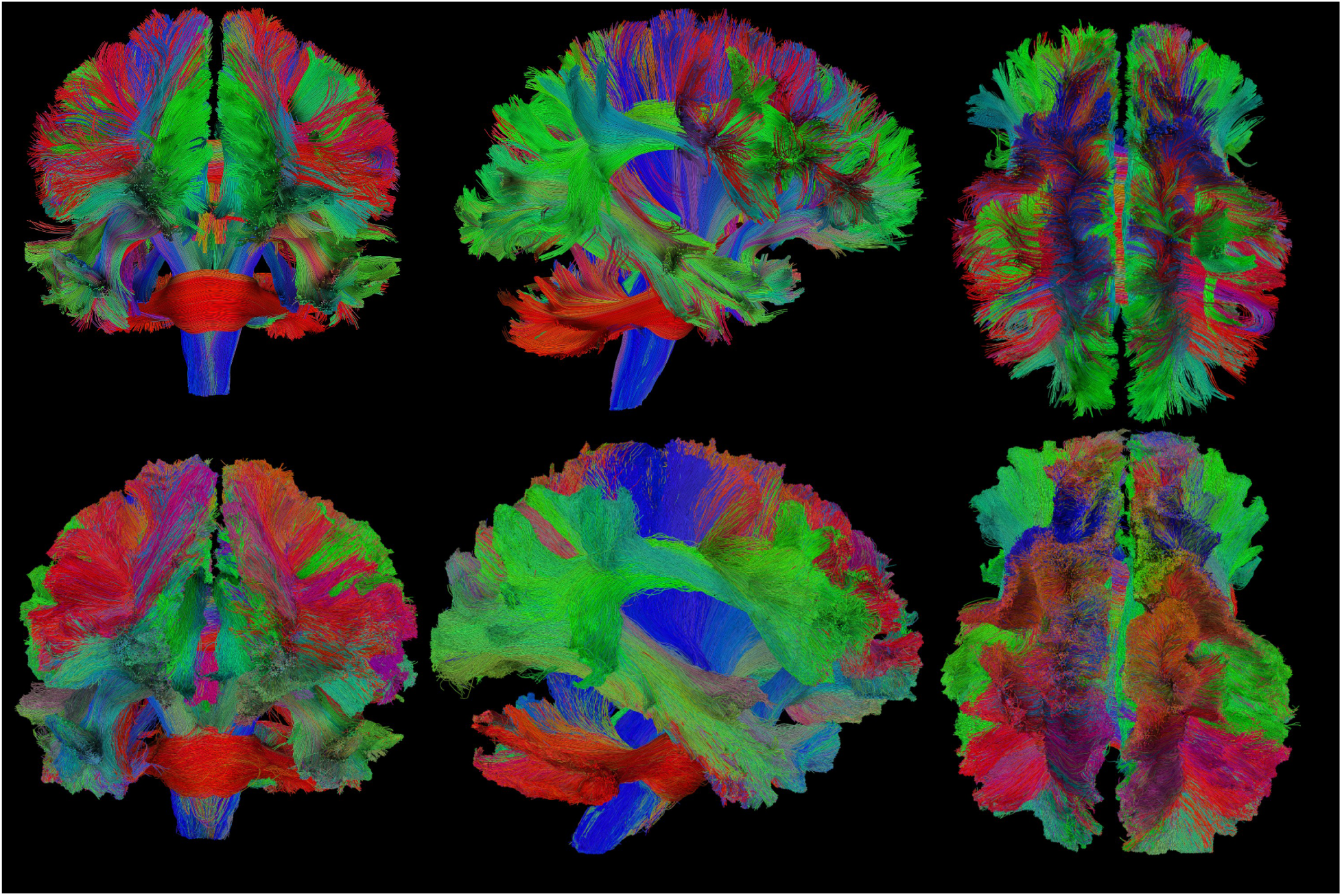
Visualization of the ground-truth ISMRM2015 tractogram and its reconstruction by the proposed method. *Top row*: Ground-truth bundles. *Bottom row*: Reconstruction through reinforcement learning.

Our method is competitive compared to prior methods without the need for labelled data. While previous ML-based method exhibit high performance when trained on the ground-truth data (which is never available on real *in-vivo* data), we can see a clear drop in their performance when trained on manually segmented tractograms (cf. rows 6-7 vs 8-9). As such, while our method is outperformed on some metrics by algorithms trained on “perfect” ground-truth data, we outperform to ML algorithms trained on realistic labelled data.

To this extend, we achieve a higher VC rate than Learn-to-Track[35], which was trained on GT data, as well Benou et al.’s DeepTract and Neher et al.’s random forest trained on realistic data. We also report higher overlap than supervised methods trained on realistic data, the lowest NC rate of all methods as well as a similar number of valid bundles as all prior supervised learning methods.

### 4.3 Experiment 3: Generalization through the HCP and ISMRM2015 datasets

In Table 3, we compare our method against Learn-to-Track [35] with both the raw diffusion signal and the fODF SH coefficients concatenated with the white-matter mask as input. We also provide results for prior methods. Overall, our SAC method outperforms TD3 as well as every other methods especially when considering the VC, IC and NC rates. We analyze the high number of invalid bundles (IB) produced by our method in section 5.4.

**Table 3:** Results obtained after training on the HCP dataset and testing on the ISMRM2015 dataset. Learn-to-Track was trained by the authors and scores for Neher et al.[31], Wegmayr et al.[59] as well as Entrack[58] were extracted from Entrack [58]. ‘X’ marks metrics that were not reported.

| | VC (% ↑) | VB (↑) | OL (% ↑) | IC (% ↓) | IB (↓) | NC (% ↓) |
| --- | --- | --- | --- | --- | --- | --- |
| Track-To-Learn (TD3) (Ours) | 53.6 | 23 | 58.0 | 45.2 | 181 | 1.1 |
| Track-To-Learn (SAC) (Ours) | 68.5 | 23 | 65.8 | 30.7 | 186 | 0.8 |
| Learn-to-Track (DWI) [35] | 27.5 | 24 | 56.2 | 55.8 | 218 | 16.7 |
| Learn-to-Track (fODF + WM Mask) [35] | 49.8 | 23 | 65.6 | 43.1 | 172 | 7.1 |
| Neher et al.[31, 32] | 52 | 23 | 59 | X | 94 | X |
| iFOD3 [59] | 72 | 23 | 16 | X | 57 | X |
| Entrack [58] | 52 | 24 | 58 | X | 123 | X |

### 4.4 Experiment 4: Impact of the discount parameter

Here we discuss the impact of the discount parameter on the (FiberCup) Tractometer metrics. We report some of the final metrics extracted by the Tractometer plotted against the chosen discount for the training runs in figure 7.

**Figure 7:**
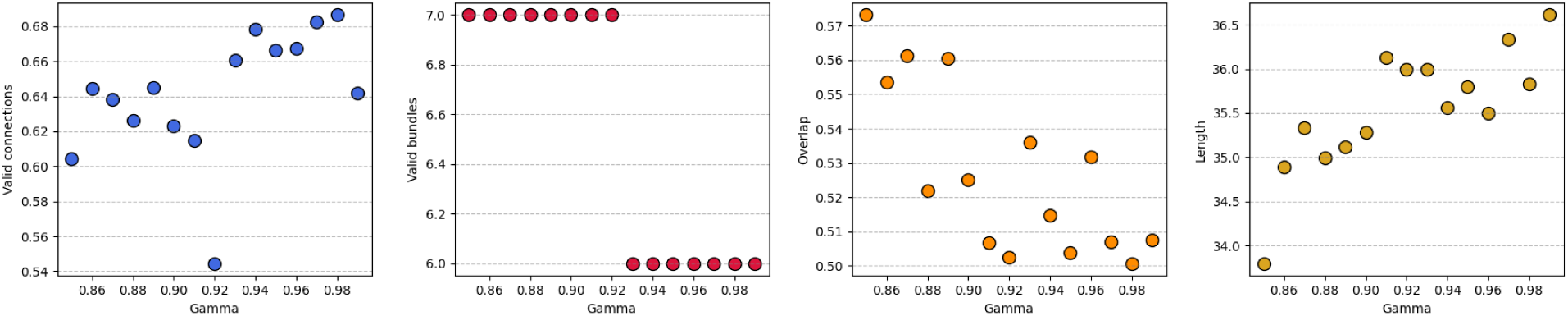
Relation between the discount (*γ*) parameter and (FiberCup) Tractometer metrics. While the method is quite robust to a wide range of discount values, there seems to be a correlation between the discount factor and the VC, the overlap and the tracks length.

As we can see, our method seems to be robust to a wide variety of discount parameters. We can observe that a higher discount generally leads to a higher VC rate in our experiments, as well as longer streamlines. It makes sense that a lower discount would lead to shorter streamlines as the agent has less incentives to keep tracking as future actions being worth less. We can also observe that a higher discount leads to reduced overlap. As such, even though no catastrophic results can be observed from a “bad” choice of *γ*, it should nonetheless be chosen with care.

### 4.5 Experiment 5: Impact of the exploration rate

We report metrics extracted by the (FiberCup) Tractometer against the chosen exploration noise in figure 8. As expected, having a higher exploration rate seems to lead towards a higher true-positive rate, more bundles reconstructed and a better overlap, but only up to a certain point. Once the *σ* parameter is set too high, the agent cannot properly exploit good actions and the overall performance of the agent decreases.

**Figure 8:**
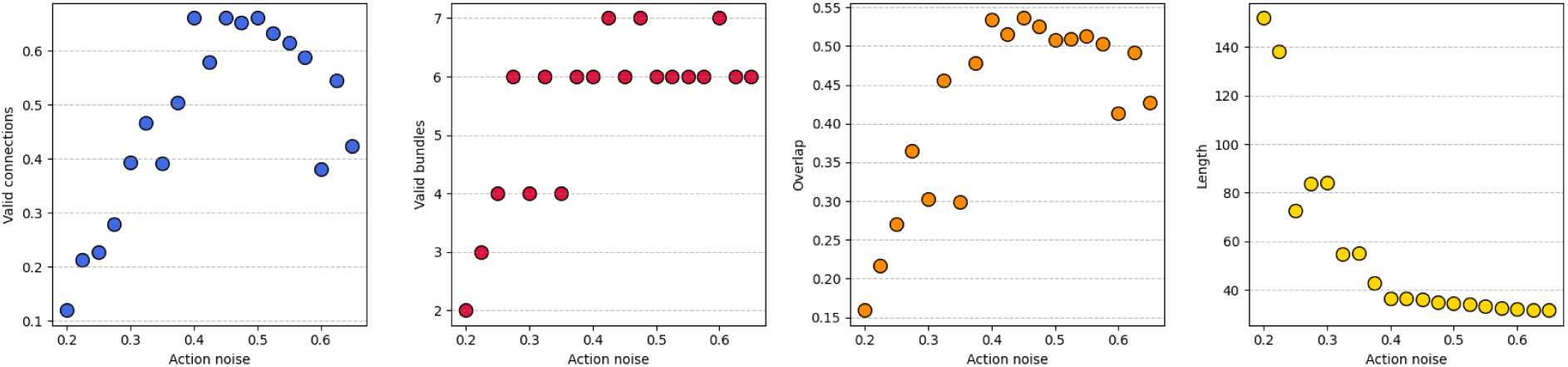
Analysis of the exploration noise versus (FiberCup) Tractometer metrics. We can see that a higher exploration noise generally leads to a higher valid connection rate, but too much noise and the performance decreases.

Similarly, too much noise added to the agent’s actions forces the agent to an erratic behaviour and more spurious actions, which may drive the streamlines out of the tracking mask prematurely, leading to a reduced mean streamline length. Yet, too little noise might allow the agent to be able to exploit the reward function and track “indefinitely”, leading to bad performance, as discussed in section 5.3. As such, the exploration rate must be carefully chosen in order to maximize performance.

### 4.6 Experiment 6: Impact of noise at test time

In table 4, we see that the level of noise added at test time allows to control the trade-off between valid connections and overlap. With no noise, the VC rate is at its highest but the OL suffers to the point where all bundles may not be reconstructed. As the added noise gets higher, the OL increases but the VC rate suffers, until too much noise is added and the overall performance degrades. While allowing the agent to control its own noise does seems appealing, we can see that the actions output by the agent are too noisy and lead to poor performance.

**Table 4:** Table reporting Tractometer metrics for the same agent with varying levels of noise added during tracking on the ISMRM2015 dataset.

| | VC (% ↑) | VB (↑) | OL (% ↑) | IC (% ↓) | IB (↓) | NC (% ↓) |
| --- | --- | --- | --- | --- | --- | --- |
| Track-To-Learn (SAC) Stochastic | 49.2 | 23 | 70.4 | 46.1 | 200 | 4.7 |
| Track-To-Learn (SAC) 0.0 | 69.1 | 22 | 51.9 | 30.4 | 147 | 0.5 |
| Track-To-Learn (SAC) 0.05 | 69.0 | 23 | 56.5 | 30.4 | 141 | 0.5 |
| Track-To-Learn (SAC) 0.1 | 68.4 | 23 | 62.1 | 30.8 | 161 | 0.8 |
| Track-To-Learn (SAC) 0.125 | 66.8 | 23 | 65.1 | 32.0 | 166 | 1.2 |
| Track-To-Learn (SAC) 0.15 | 63.4 | 23 | 67.8 | 34.1 | 175 | 2.4 |
| Track-To-Learn (SAC) 0.175 | 58.0 | 23 | 69.9 | 36.7 | 178 | 5.3 |
| Track-To-Learn (SAC) 0.2 | 50.6 | 24 | 70.6 | 39.4 | 170 | 10.0 |
| Track-To-Learn (SAC) 0.225 | 42.7 | 23 | 68.2 | 41.5 | 137 | 15.8 |
| Track-To-Learn (SAC) 0.25 | 35.4 | 23 | 61.9 | 42.9 | 121 | 21.8 |

## 5 Discussion

### 5.1 Performance

Experiment 1 and 2 show us that despite a simple reward function and architecture, the proposed RL method is able to achieve highly-competitive performance when compared against classical and supervised-learning based methods. From the results presented in sections 4.1 and 4.2, we can appreciate a high VC rate, high number of reconstructed VB, a decent overlap and a very low rate of NCs.

Looking back at the implementation, and comparing the results against classical methods, we can interpret the Track-To-Learn framework as the groundwork for data-driven improvements over classical algorithms. While the reward function allows the learned agents to behave similarly to classical methods, the controllable level of noise allows the user tracking with Track-to-Learn to directly influence the resulting tractograms. It allows the user to fine-tune the trade-off between probabilistic and classical algorithms, instead of having to choose between the two.

Having a low rate of NCs is very interesting: no-connections indicate streamlines that do not connect two regions. Anatomically speaking, no fibers in the white-matter can have this configuration. As such, these streamlines are usually discarded when performing connectivity analyses, for example, because they most-certainly do not represent the underlying anatomy. While prior methods like Neher et al.[32] had to, for example, implement a hard prior to have streamlines “bouncing off” the white-matter mask to prevent it from terminating prematurely, our agents learned to diverge the tracking process from exiting the white-matter mask on their own.

### 5.2 Generalization capabilities

One of the stronger aspects of the proposed method is certainly its generalization abilities. From experiment 1 and 3, we can observe really strong performance when dealing with a different dataset than the one used for training. Notably, let’s take a look at Experiment 1: notice the dip in performance when the prior supervised-learning method tracks on the “flipped” dataset. By looking at figure 5, we can see that parts of the reconstructed tractogram seem to try to replicate the original FiberCup instead of reconstructing new, never-seen fibers as would a classical algorithm do. For example, we can see that the streamlines in bundle 7 try to “go back” and follow the direction that would have been correct in the original FiberCup tractogram. Meanwhile, agents trained using the Track-To-Learn framework reconstruct very similar tractograms. Even though the resulting streamlines are noisier, we can see that all bundles are reconstructed, and that “quirks” learned on the FiberCup dataset are replicated in the flipped version.

Experiment 3 not only presents very strong results when training on the HCP dataset, but also reinforces the notion that the algorithm generalizes to other datasets. Indeed, comparing the performance presented between experiment 1 and 3, we can notice a hard drop in performance from other ML methods like Neher et al.[31] and Poulin et al.[35]. We report similar performances between the two experiments, implying that our generalization capabilities are not upper bounded by the change in datasets, but rather by the reconstruction capabilities of the proposed method, which could be increased in future works.

### 5.3 Analysis of reward function

As mentioned before, a good reward function encompasses the underlying goal of the agent. As such, if an agent maximizes its cumulative reward, it should produce increasingly accurate results. To see if the reward function introduced in section 2.2.3 truly satisfies our goal of producing good tractograms, we analyzed the sum of rewards obtained during the hyperparameter search for experiment 1. To do so, we plotted the sum of rewards obtained by TD3 against the VC ratio and OL produced by the Tractometer during their last validation run. Because longer trajectories may indicate more reward (as the agent receives a reward during more timesteps), we also plotted the length of the produced streamlines for the same validation runs.

Scatter plots in figure 9 indicate that achieving a higher cumulative reward does mean better performance, but only up to a certain point. We can also see a strong correlation between the reward obtained and the length of the streamlines.

**Figure 9:**
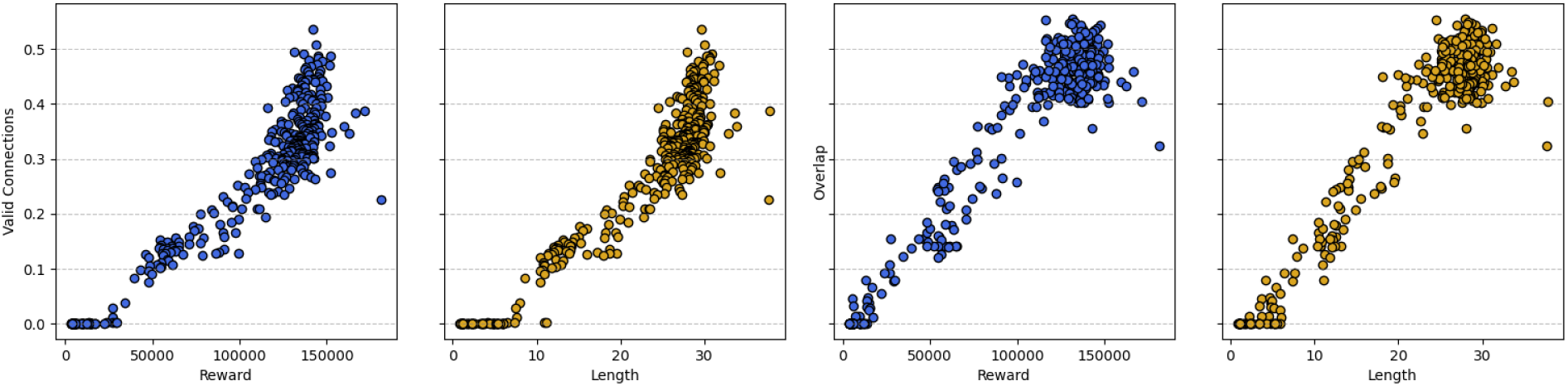
Analysis of the relation between VC and OL (y-axis) versus the final reward obtained by the agent and the mean length (in mm) of the streamlines produced during validation (x-axis). Results come from an hyper-parameter search for Experiment 2. We can see that a higher reward or mean length generally means a higher VC rate and OL up to a point. We also observe a strong correlation between the total reward obtained and the length of the produced streamlines.

To see why performance degrades when the reward is too high, we can take a look at an agent achieving very high reward on the FiberCup. Figure 10 shows training curves extracted from an agent achieving high reward. Around the 80th episode, the VC rate drops as the accumulated reward and streamline length blow up. As a comparison, the mean streamline length in the FiberCup’s ground-truth set of streamlines is 114mm and the maximum length is of 145mm. Therefore, a final mean length of *∼*150mm (3rd plot) is too much. A visual inspection of the reconstructed streamlines, available in appendix A, reveals that the agent is able to circumvent streamline termination by “looping”, allowing it to track for much longer than it should, therefore accumulating more reward.

**Figure 10:**
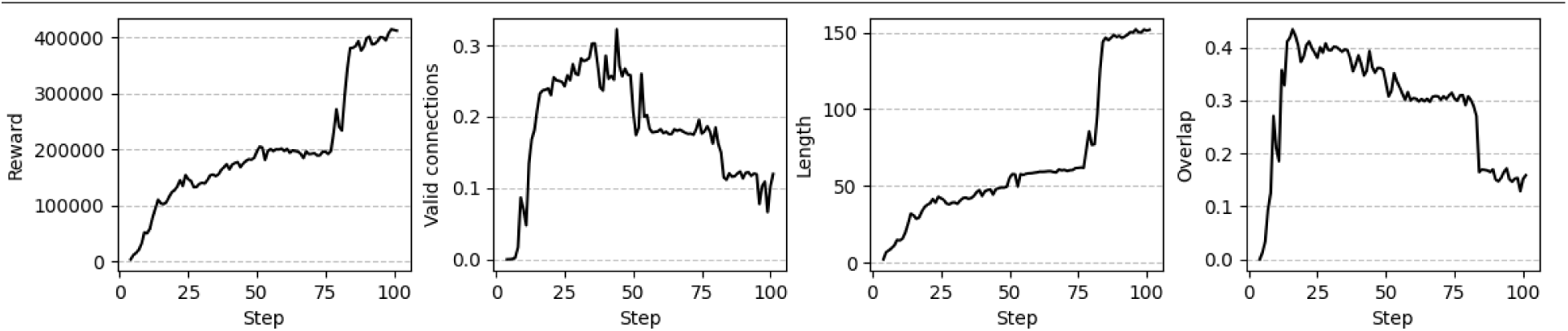
Training curves from an agent achieving high reward on the FiberCup dataset. We can see that a jump in accumulated reward and streamline length around the 80th episode comes with a dip in measured VC rate and OL.

Reward hacking [1,13] happens when the agent finds a way to maximize its accumulated reward while violating the overarching objectives it was supposed to reach. For example, an agent might find a bug in a video game allowing it to achieve a high score without actually progressing in the game and therefore demonstrating knowledge and comprehension of it. By examining the streamlines in appendix A, we can observe that the reward function introduced in section 2.2.3 and the episode-termination conditions introduced in 2.2.1 still allow the agent to gather high reward without producing meaningful tractograms. As such, while the proposed reward function does lead to competitive performance, careful monitoring is still required to make sure it is not taken advantage of by learning agents.

### 5.4 Analysis of invalid bundles reconstructed

Despite good overall results, experiments 2 and 3 display a large number of invalid bundles. Taking into account the fact that the proposed method works by exploring its environment and that global connectivity, in the form of bundles, must be reconstructed from local information, the fact that the proposed method reconstructs a lot of invalid bundles is not very surprising. Here we analyze how the invalid bundles reconstructed by the SAC agent compare against its valid bundles. Figure 11 provides some statistics on the reconstructed bundles.

**Figure 11:**
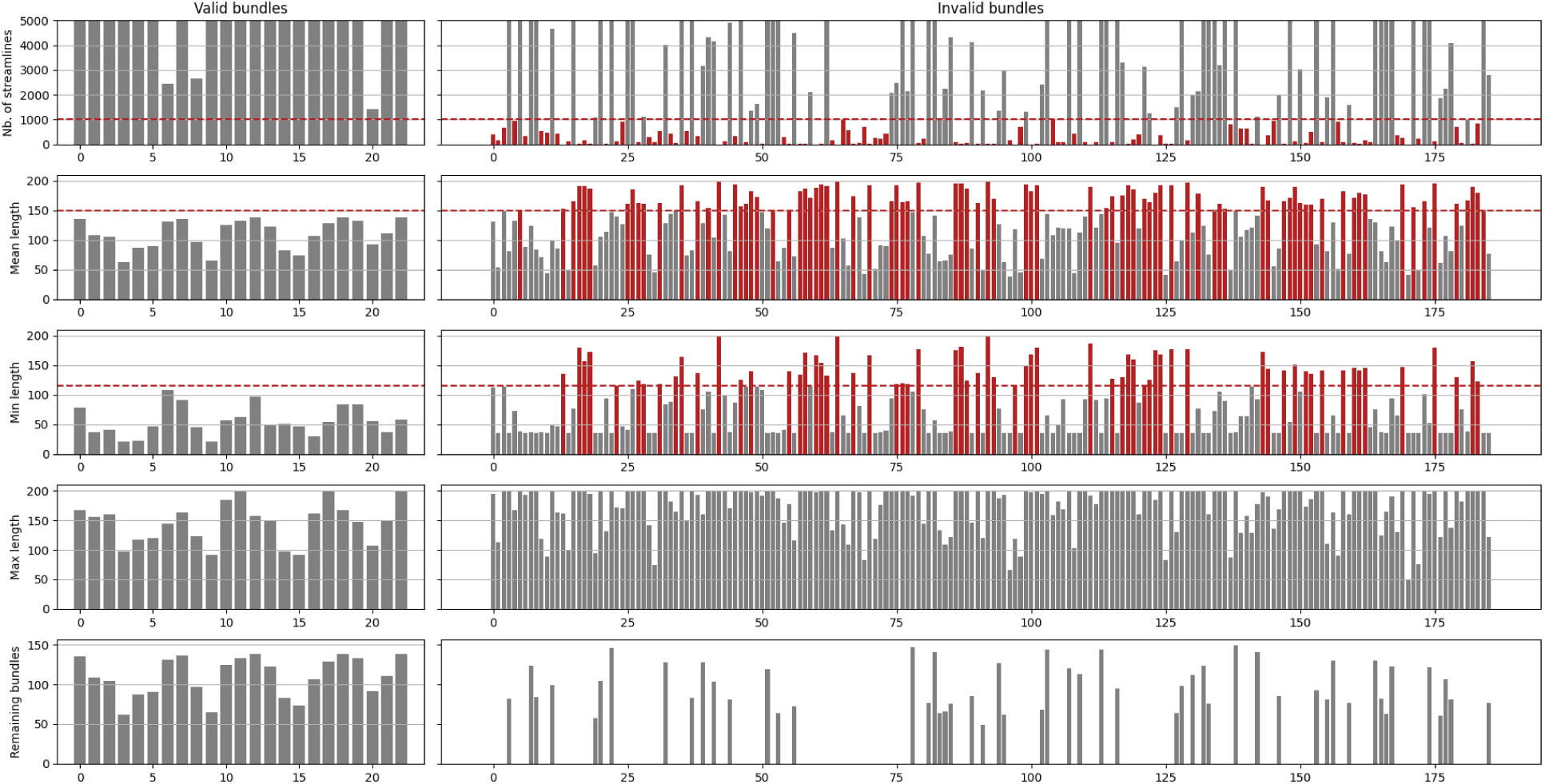
Measures of valid and invalid bundles reconstructed in Experiment 3 by the SAC agent. Y-axis is shared across columns. Red bundles could be discarded according to simple criteria. Left column: measures of valid bundles. Right: measures of invalid bundles. First row: Number of streamlines per bundles, where the y-axis has been cut-off for clarity. Second row: Mean streamline lengths per bundle reconstructed. Third row: Minimum streamline length per bundle reconstructed. Fourth row: Maximum streamline length per bundle reconstructed. Fourth row: Mean length of the 52 (out of 186) remaining bundles after pruning via simple criteria.

**Figure 12:**
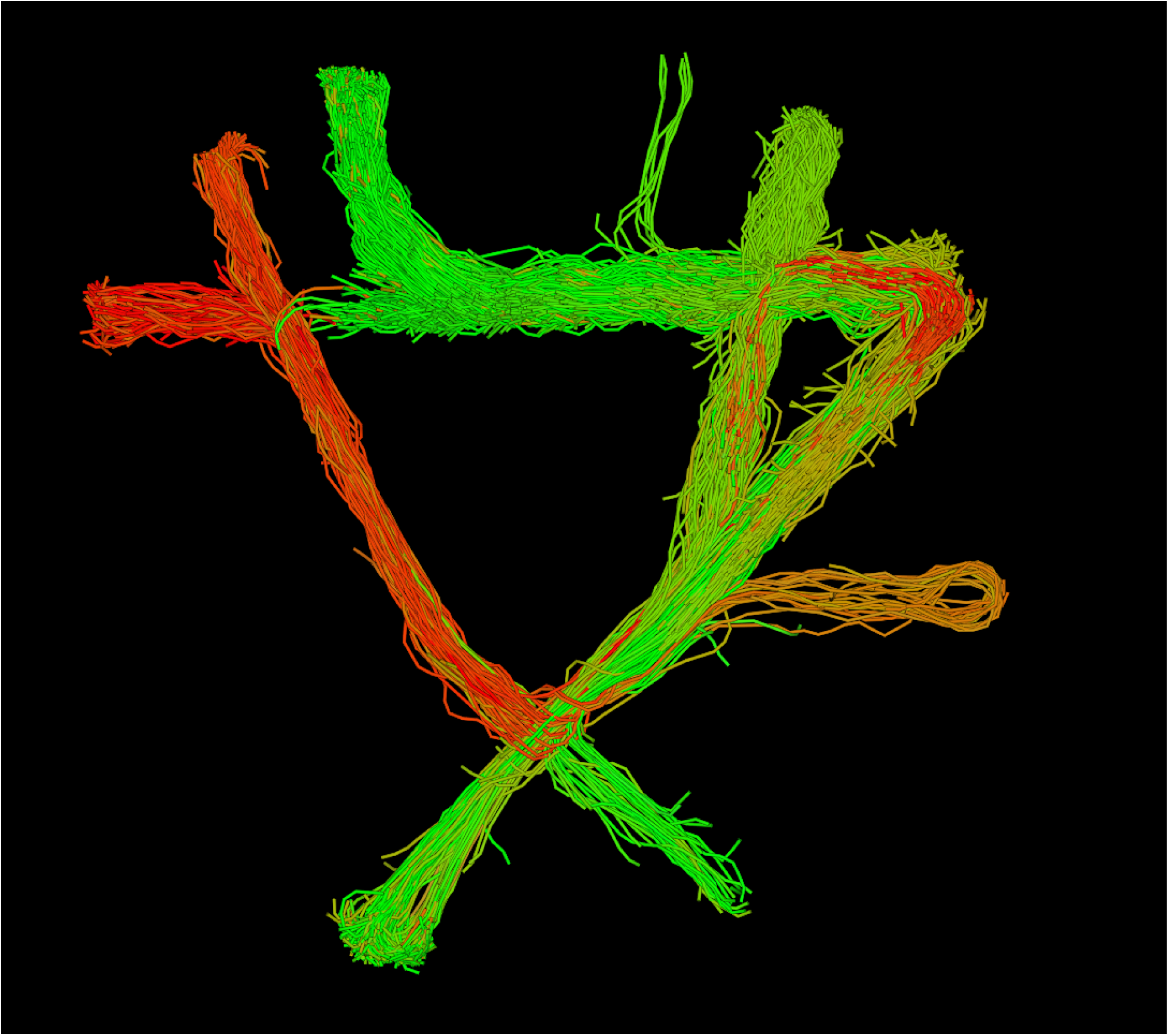
Tractogram reconstructed on the FiberCup phantom, with the shortest streamlines removed. As we can observe, even with the constraints listed in section 2.2.1, the agent might still exploit the environment to track for much longer than it should, performing loops instead of terminating.

The first row of figure 11 indicates that the invalid bundles reconstructed by our agents tend to have a very low streamline count. From row 2 and 3, we can observe that invalid bundles tend to have either very long or very short streamlines. Therefore, while a gross filtering of bundles based on their streamline length cannot be considered as it would lead to the elimination of valid bundles, further post-processing of bundles could be done to reduce the number of false-positive. For example, no valid bundle has a mean streamline length above 150mm or a minimum streamline length over 115mm. Row 5 displays the mean length of the 52 (out of 186) bundles that would remain if we were to perform filtering based on the aforementioned criteria.

Therefore, while agents trained via the proposed framework do tend to reconstruct many invalid bundles, we can observe that invalid bundles tend to expose very different characteristics from their valid counterparts and that simple post-processing via bundling and thresholding of the reconstructed streamlines [16] could be used to reduce the number of IBs.

### 5.5 TD3 vs. SAC

While both RL agents achieve highly competitive results, the SAC agent seems to often outperform TD3.

The TD3 and SAC algorithms share many similarities: both employ two critics predicting a q-value for the actions output by their respective actors. In our implementation, both agents use neural networks with the same number of layers and the same width per layer. As such, the main difference comes in the form the policy takes for the algorithms. TD3 employs a deterministic policy, where its output is a vector of three components representing the action. SAC, however, employs a stochastic policy where its output is the mean and standard deviation of a 3D Gaussian distribution, from which actions can be sampled. SAC’s policy can be made deterministic by using the predicted mean of the distribution as the actions directly.

Therefore, both algorithms employ different strategies for exploration. As mentioned in section 3.3, TD3 requires explicit noise to be added to the actions output by its deterministic policy. On the opposite, SAC can handle its own exploration by sampling its policy.

We believe it is fair to draw comparisons between the two RL algorithms and classical deterministic and probabilistic algorithms: TD3 outputs direct actions as would a deterministic algorithm, while SAC tries to model the input noise as would a probabilistic algorithm. Adding further to similarities, SAC employs a lower maximum angle between streamline segments than TD3, like would probabilistic algorithms compared to deterministic ones. We determined the maximum angle for each algorithm empirically and found that TD3 would suffer from slow and poor learning with a lower maximum angle, while SAC would produce too many invalid connections and bundles with a higher maximum angle.

### 5.6 Alternatives to feed-forward neural networks

Initially, and motivated by Poulin et al.[35], we attempted to use recurrent neural-networks instead of their feed-forward counterpart. TD3 uses a replay buffer to store transitions and reuse them for training and so the replay buffer was adapted to also store the memory of the network when it produced the transition. However, with this setup we found the network to be incapable of learning the “directionality” of the streamline. Indeed, the network would learn to map input states to a single action, making going “forward” during tracking possible, but going “backwards” impossible as the network would produce actions as if it was going the other way. As such, recurrent networks were dropped in favour of simple feed-forward networks.

While the exact reason for this incapability of learning the directionality of tracking remains unknown, we made several hypothesis as to why this is the case. First, the effects of having two environments instead of one as is usual in most RL problems are still yet to be investigated. Because the network has to “re-track” the flipped half-streamlines and new predictions are replaced with the pre-existing segments, the memory of the network might no longer represent a meaningful representation of the signal, preventing the learning process from differencing different directions. Also contrary to other projects where RNNs were used in conjunction with RL [3,4,25], only a single learning agent was used instead of multiple. Finally, Kapturowski et al.[25] proposes to have the network predict a couple of “burn-in” steps before truly learning on the transitions to fight saved memory from going “stale”, which was not done in our case.

### 5.7 Future works

Distancing ourselves from prior work, we chose to feed the agent the spherical harmonics coefficients from fiber ODFs instead of the (possibly resampled) raw diffusion signal. This imposes a prior on the data which previous machine learning methods for tractography generally did not have. As such, the proposed method should also be tested using the raw resampled diffusion signal as input, to provide a fairer comparison.

Having two environments makes the present method harder to adapt for reinforcement learning algorithms. Many of them require asynchronous actors learning and having two environments makes this process rather awkward. Therefore, the process of tractography could be adapted to only require one environment by exclusively seeding from the grey-matter/white-matter interface and thus removing the need to go “backwards” as did Wegmay et al.[59]

As discussed in section 5.6, unsuccessful attempts were made to take advantage of the memorial capacities of recurrent neural networks. While appending the last four directions of the streamlines is one way to correct for the directional dependencies of tracks, we believe that more work needs to be done so that RNNs can really be successfully used for tractography.

While our method shares similarities with probabilistic tracking methods, our agents tend to produce “skinnier” bundles with a lower OL than their classical counterparts, as well as a lower overlap than some prior ML methods. While the reasons for this require more investigation, we can speculate that the agents tend to stay away from the limits of the white-matter mask in fear of terminating. Since the agent is incentivized to track for as long as possible so as to receive a large reward, tracking near the limits of the white-matter is riskier, potentially leading to a lower OL. To alleviate this problem, probabilistic termination maps [18] could be used instead of a binary WM mask to determine streamline termination.

As mentioned in section 5.4, the proposed method tends to reconstruct a high number of invalid bundles, as per its exploratory nature and the fact that, reward-wise, invalid bundles are worth as much as valid ones. As such, the reward function must be improved to allow better disentanglement between valid and invalid bundles, probably with an atlas or shape-priors. Because designing a reward function by hand is difficult, an alternative would be to learn the reward function through *inverse reinforcement learning*, where both the policy and the reward function are learned at the same time.

## 6 Conclusion

The ill-posedness of the tractography inverse problem suggests, in our opinion, that reinforcement learning methods are best suited to solve it. RL methods can exploit the expressive capabilities of neural networks without the need for hard-to-obtain and/or biased labelled-data. Therefore, we presented the deep reinforcement learning *Track-to-Learn* framework. The framework allows for data-driven improvements over classical methods by making a learning agent improve its tracking abilities iteratively via trial-and-error. We report highly competitive results on a variety of datasets. While prior supervised learning methods struggled with generalization to new, unseen datasets, we report little to no loss of performance when tracking on unknown diffusion volumes. Even with the reported results, the presented method lays the groundwork for tractography via reinforcement learning. There is much room for improvements and our open-source Track-to-Learn framework can used to build, in the future, even better tractography algorithms.

## Acknowledgments

The authors would like to thank members of the SCIL^8^ and VITAL^9^ groups for their suggestions, insight and discussions on this project.

## Footnotes

2 Code will be rendered public upon acceptance of this paper.

3 By comparison to their feed-forward alternative, RNNs produce an internal representation of the input data alongside each prediction. This internal representation is then fed to the next input so that it can be reused, allowing the network to build a memory of past inputs and predictions.

4 Diffusion MRI signal being symmetric, peaks can be followed in any direction.

5 https://comet.ml

6 https://github.com/scilus/scilpy

7 https://github.com/scilus/scilpy

8 http://scil.dinf.usherbrooke.ca/en/

9 http://vital.dinf.usherbrooke.ca/

## Notes

### Competing Interest Statement

The authors have declared no competing interest.

